# Spatial context of trait variation: morphology of spotted salamanders (*Ambystoma maculatum*) varies more within ponds than between ponds

**DOI:** 10.1101/2020.06.15.153312

**Authors:** Elizabeth T. Green, Anthony I. Dell, John A. Crawford, Elizabeth G. Biro, David R. Daversa

**Author notes:** Corresponding Author: Dave Daversa.

## Abstract

The influence of intraspecific trait variation on species interactions makes trait-based approaches critical to understanding eco-evolutionary processes. Because species occupy habitats that are patchily distributed in space, advancement in trait-based ecology hinges on understanding how trait variation is distributed within and between habitat patches. We sampled larval spotted salamanders (*Ambystoma maculatum*) across spatially discrete ponds to quantify within- and between-pond variation in mass, length, and their allometric relationship. Between-pond variation explained 7-35% of total observed variation in the length and shape of salamander larvae, depending on the body segment measured (i.e., head, body, or tail). Salamander tail morphology was more variable and exhibited more between-pond variation than head or body morphology. Salamander mass was highly variable and strongly correlated with total length. Allometric analysis revealed that the slopes of mass-length relationships were similar across ponds, but that intercepts differed across ponds. Preliminary evidence hinted that newly constructed ponds were a driver of the observed differences in mass-length relationships. Pond construction may therefore bolster trait diversity across the broader landscape, and in so doing instil more adaptive potential of salamander populations under current and future environmental change.

---

Morphological traits underpin many eco-evolutionary processes. The range of morphology exhibited by individuals of a species, a form of intraspecific trait variation (Bolnick et al., 2011), shapes the niche breadth of populations, which in turn affects their resiliency to environmental disturbances and biological invasions (Tack et al., 2014). Because of this, models of population and community dynamics that incorporate intraspecific trait variation have become central to ecology and evolution (Bolnick et al., 2011; Moran et al., 2016). A key outstanding challenge for this ‘trait-based’ paradigm is to understand trait variation in a spatially explicit context (Violle et al., 2012; Moran et al., 2016).

Species tend to occur in landscapes comprised of spatially discrete habitat patches, and traits may vary both among individuals within habitat patches and among groups across habitat patches. Theory predicts that the influence of trait variation on population and community dynamics depends on how variation is partitioned within versus between habitat patches (Moran et al., 2016; Banitz, 2019). Between-patch variation potentially allows for a broader range of adaptive responses to environmental disturbances (Moran et al., 2016) and antagonistic interactions (Tack et al., 2014) than does within-patch variation. Alternatively, between-patch trait variation may heighten extinction risk by geographically isolating trait diversity to specific habitat patches. Population and community dynamics are influenced not just by the degree of intraspecific trait variation but also the relative proportion of trait variation that occurs within-versus between-patches (Violle et al., 2012; Moran et al., 2016).

Trait co-variation describes relationships between multiple individual traits. Between-patch variation in how traits co-vary may also be influential in eco-evolutionary processes (but see Evangelista et al., 2019). Measures of trait co-variation are useful for describing species growth patterns (Hirst et al., 2014), their adaptive constraints (Voje et al., 2014), and complex morphological characteristics such as body shape (Laughlin and Messier, 2015). Practically, they provide a tool for filling data gaps in ecological models (Madin et al., 2016). Allometric scaling of mass with body length is a basic form of trait co-variation that is widely used for this purpose (Madin et al., 2016). Additionally. morphometrics integrates co-variation between body length, depth, and sometimes width, to characterize body shapes of individuals. Body shape, being a strong proxy of performance and fitness, is arguably a better predictor of species adaptive responses to environmental change than linear body measurements (Laughlin and Messier, 2015). As such, shape and allometry can be powerful predictors of ecology and evolution in patchy landscapes.

In this study, we assessed within- and between-patch trait variation and co-variation in larval spotted salamanders (*Ambystoma maculatum*). We sought to quantify the extent of between-patch differences in salamander mass and length, as well as allometric and morphometric relationships among multiple traits. We sampled larval spotted salamanders among a network of spatially discrete wetlands (i.e., ponds) and measured body length and mass of 519 individuals. We used these data to quantify individual variation in mass and length within and among ponds. We then examined the contribution of between-patch (i.e., between-pond) differences to the total observed variation and co-variation of those traits. As an initial exploration into potential drivers of between-pond differences in salamander morphology, we also performed a preliminary examination of whether salamander mass, length, mass-length allometry, and shape were influenced by the age and predator density of ponds.

## Materials and Methods

### Study species

Spotted salamanders are broadly distributed throughout the Northeastern and Midwestern United States (Petranka, 1998). Spotted salamanders are semi-terrestrial pond breeders, annually migrating from terrestrial hibernacula to reproduce in fishless wetlands. Breeding in our study area occurs between March-April (Sexton et al., 1990). After hatching from eggs, aquatic larvae develop and metamorphose in 6-10 weeks (Petranka, 1998). Larvae feed on invertebrates and anuran tadpoles, and are themselves prey for adult salamanders and larger aquatic invertebrates such as odonate larvae and beetles (Urban, 2010). Spotted salamander populations are useful systems to study morphological variation in a spatial context because: i) individuals occupy and move among spatially discrete ponds that comprise functional metapopulations with patchy habitat structure (Patrick et al., 2008); and ii) larval stages exhibit substantial morphological plasticity in response to heterogeneity in biotic and abiotic conditions (Scott, 1990; Urban, 2010; Shaffery and Relyea, 2015), which permits a range of trait expressions across individuals and ponds.

### Field sampling and husbandry

We sampled multiple ponds in east-central Missouri, distributed across three distinct conservation properties - Tyson Research Center (800 ha), Forest 44 Conservation Area (400 ha) and Shaw Nature Reserve (700 ha) (Fig. 1). We focused on six ponds in which pilot surveys confirmed Spotted salamander larvae were present. Three ponds (Mincke Pond, Arthur Christ Pond, Beth’s Pond) were constructed in 2008 for research purposes and had similar sizes and dimensions (Burgett 2015). As part of a separate experiment, Rotenone was applied to Beth’s Pond in 2008. While this initially reduced microbial biodiversity (Woods et al., 2016) the micro-organismal community structure had returned to pre-treatment conditions well before sampling for this study (Woods et al., 2016). The other three focal ponds (Salamander Pond, Forest 44 Pond, Shaw Pond) were older and variable in size (Table S1). Salamander Pond was created in 1965, and Forest 44 Pond and Shaw Pond between 1990 and 1996 (data extracted from Google Earth Historical Imagery). All ponds contained multiple predators of larval spotted salamanders including: dragonfly (*Anax* sp.) and damselfly (*Lestes* sp.) nymphs, diving beetles (*Dytiscidae* sp.), hyrdrophylid beetles (*Tropisternus* sp), and adult newts (*Notophthalmus viridescens*) (Tables S1 and S4; E.G. Biro unpublished data). All ponds were located within forested habitats typical of temperate deciduous ecosystems found in the Midwestern US. A major highway bisected the study area, separating Forest 44 and Shaw ponds from the other four ponds.

**Fig. 1.**
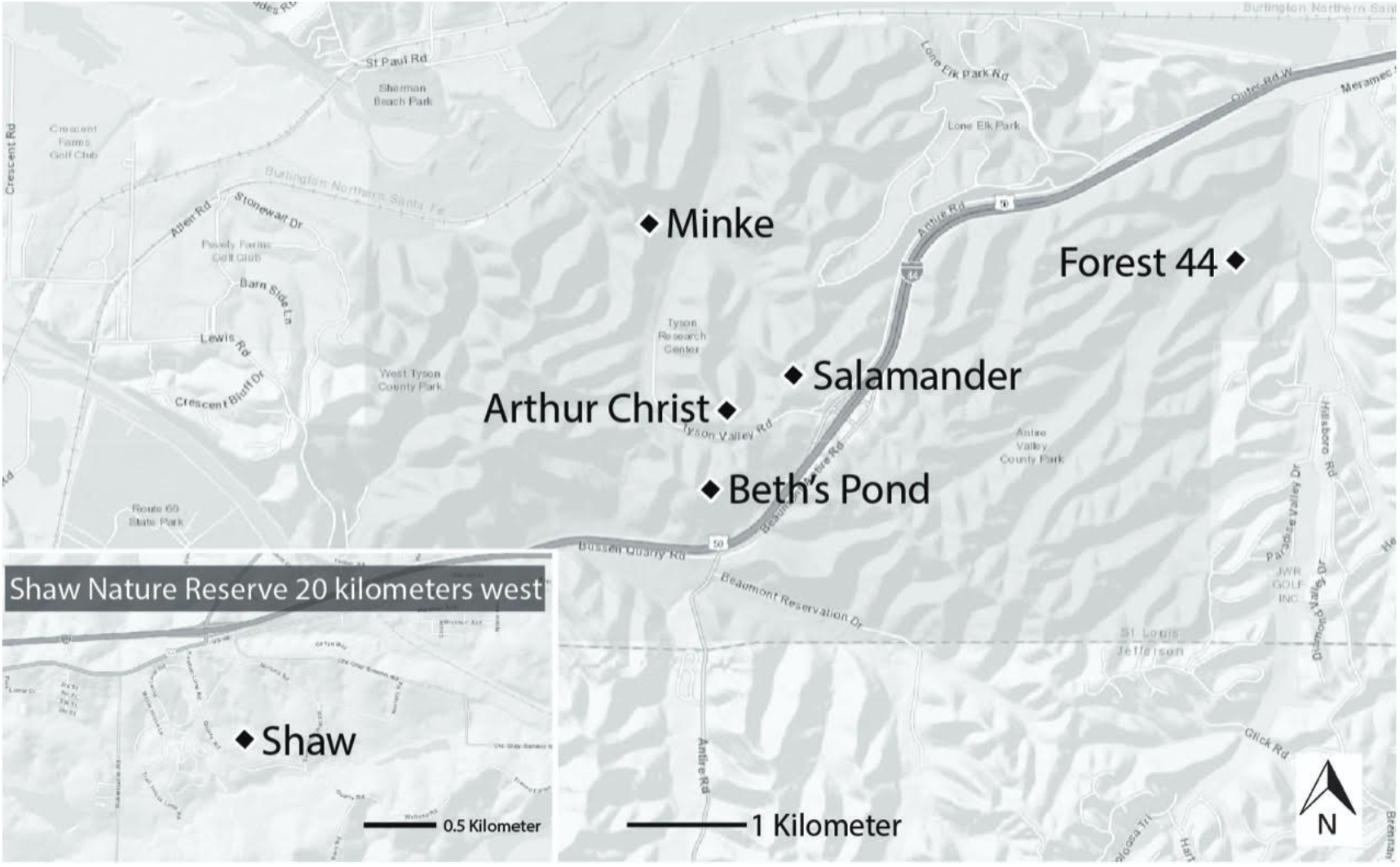
Map of study area. The six focal ponds were located in Eastern Missouri (US across three conservation areas. Mincke Pond, Arthur Christ Pond, and Beth’s Pond are located in Tyson Research Center. Shaw Pond is located in the Shaw Nature Reserve. Forest 44 Pond is located in Forest 44 Conservation Area. All ponds occurred in Oak-Hickory forests typical of the region.

We intensively sampled one pond per week (30 June-08 August 2016) by dip-netting near the perimeter of the pond. This sampling design confounded between-pond variation in spotted salamander morphology with possible temporal morphological variation due to growth and development (Landberg and Azizi, 2010; Musseau et al., 2020). To minimize the influence of temporal factors in salamander morphological variation, we focused sampling on the latter stage of the developmental period of salamanders (Harrison stage 45-46; Harrison, 1969), when growth and development had slowed (Landberg and Azizi, 2010). At each sampling event, we dip-netted until 10 minutes passed without a capture to maximize coverage of morphological variation within ponds. We retained all larvae that did not show overt signs of injury or illness (e.g., damaged tails or legs, tumorous growth, etc.) and immediately transported them to the National Great Rivers Research and Education Center (NGRREC) - less than 1 hr drive - where they were housed for seven days before being returned to their original ponds.

We housed larvae individually in circular plastic arenas (28 cm diameter) filled with 500 mL dechlorinated water (approximately 2.5 cm depth). Larvae were maintained at 18°C with a 14:10 h light:dark cycle, consistent with ambient conditions at the surveyed ponds. Salamanders were fed a single gray tree frog (*Hyla chrysoscelis-versicolor*) tadpole on the fifth day as part of a separate experiment. Observing that not all salamanders ate the tree frogs fed to them the prior day, we tested whether feeding influenced measures of mass for a random subset of 237 individuals for which feeding data were available (see Supplementary Material for detailed methods).

As an initial exploration into potential drivers of between-pond variation in salamander morphology, we assessed whether salamander mass, length, mass-length co-variation, and shape were influenced by the age and predator density of ponds. We consider these assessments preliminary because of the low replication of ponds in our sample (N = 6). We used historical information described above to classify pond age as ‘new’ (N = 3; Mincke Pond, Arthur Christ Pond, Beth’s Pond) or ‘old’ (N = 3; Salamander Pond, Forest 44 Pond, Shaw Pond) (Table S1).

To estimate predator density, we systematically dip-netted the focal ponds and recorded the abundance and composition of two broad types of predators of spotted salamander larvae: macro-invertebrates and adult amphibians (Table S5, below); none of the focal ponds contained fish. We were unable to sample Beth’s Pond for predators, and instead used historical data collected by E.G.B. in 2013. We checked predator density counts in 2013 against our 2016 sampling using Mincke Pond, the pond for which we had data from both years. Predator densities in Mincke Pond in 2013 were similar to those that we observed in 2016, so we considered our predator density estimates for Beth’s Pond to be representative for our sampling period.

### Trait measurements

On the sixth day after capture, we measured the length and mass of salamanders, distinguishing between head length, body length, and tail length. We photographed lateral and dorsal images of each larvae placed into clear tanks that minimized movement (Fig. S1). We blot-dried individuals on paper towels before weighing. We measured the length of salamander heads, bodies, and tails from images using ImageJ (Fig. S1) (Rasband, 1997).

To obtain measurements of the shape of larvae we digitized landmarks on lateral images using the software *tpsDig2* (Rohlf, 2006). Following Van Buskirk & Schmidt, (2000) we tagged twenty landmarks that outlined larval shape (Fig. S1). Landmarks 1-3 described the shape of the head of the larvae, landmarks 4-11 described body shape, and landmarks 9-20 outlined tail shape. Landmarks were rotated, scaled by size, and aligned to a coordinated system using the Procrustes least-squares superimposition available in the *geomorph* package for R statistical software (Adams and Otárola-Castillo 2013). We conducted four principal component analyses to explore the scaled two-dimensional shape variation, again distinguishing head shape, body shape, and tail shape. The first principal component (PC) score accounted for most of the variation in head shape (37%), body shape (33%), and tail shape (33%). We therefore used PC1 as the shape metric in our analyses.

### Data Analyses

We calculated the coefficient of variation - the ratio of the standard deviation to the mean - as a standardized measure of individual morphological variation within ponds. We partitioned observed variation in salamander length and mass to within- and between-pond trait differences with generalized linear mixed models (GLMM), using the *Ime4* package in R (R Core Team, 2019). Specifically, we calculated the intra-class correlation coefficient (ICC) from GLMMs that included ‘pond’ as a random intercept term (Nakagawa and Schielzeth, 2010). The ICC, also called the variance partitioning coefficient (Messier et al., 2010), is the proportion of total variation in response variables that is attributable to group-level (between-pond in our case) differences. In our analysis, the ICC indicated the proportion of total observed variation in salamander length and mass that came from between-pond differences in those traits. We ran separate GLMMs for mass and the four length measurements - head length, body length, tail length, and total length. All models used a Gaussian error structure, as all morphological data were normally distributed (Fig. 3).

**Fig. 2.**
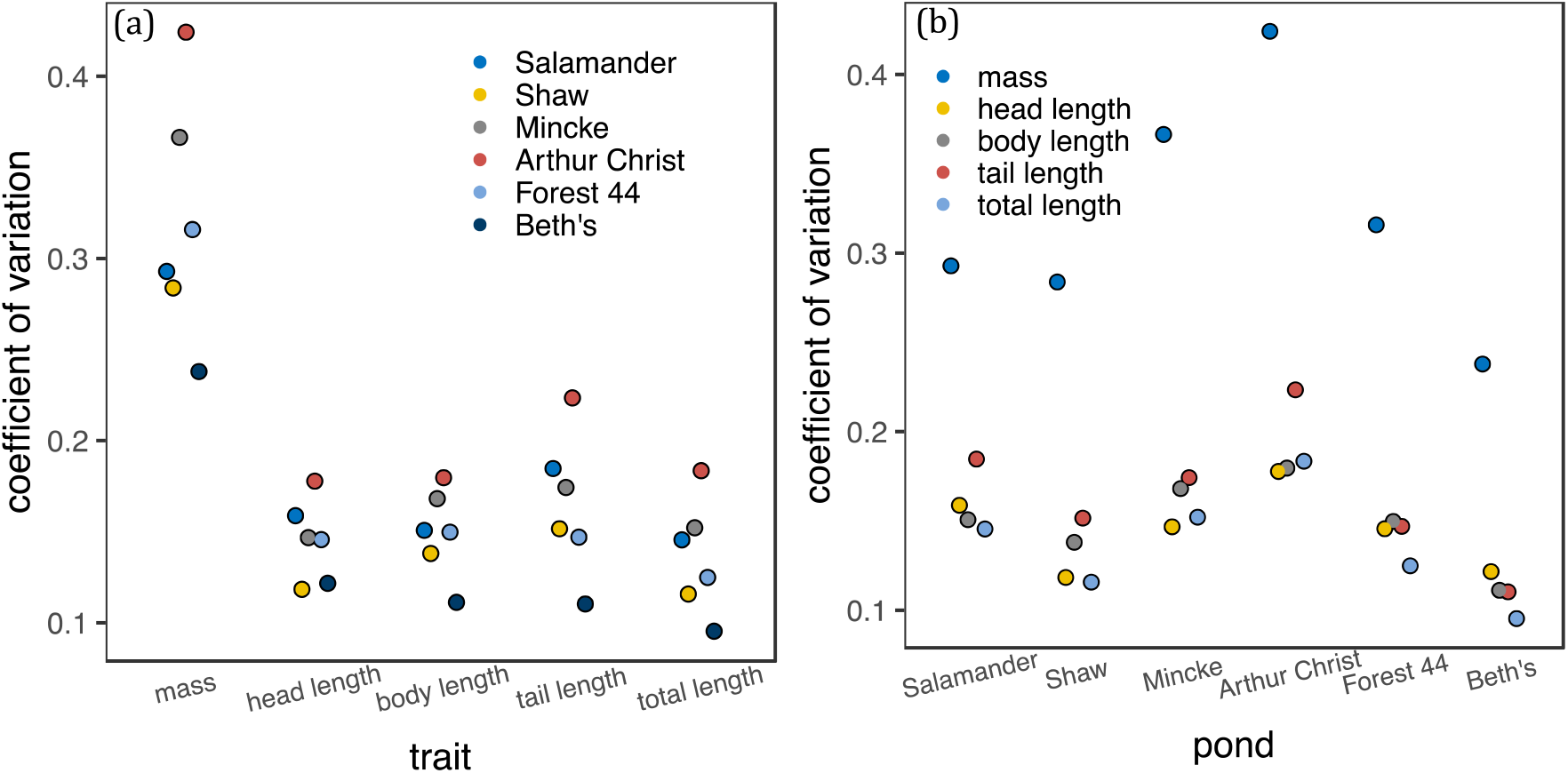
Within-pond variation in salamander morphology. The coefficient of variation, or the extent of variance in relation to mean trait values, is shown for **(a)** mass and length measures and **(b)** the six ponds where we sampled salamanders, ordered from left to right in the chronological order in which they were sampled. Colors distinguish pond of capture in (a) and the focal trait in (b). Note that the same data are reported in (a) and (b).

**Fig. 3.**
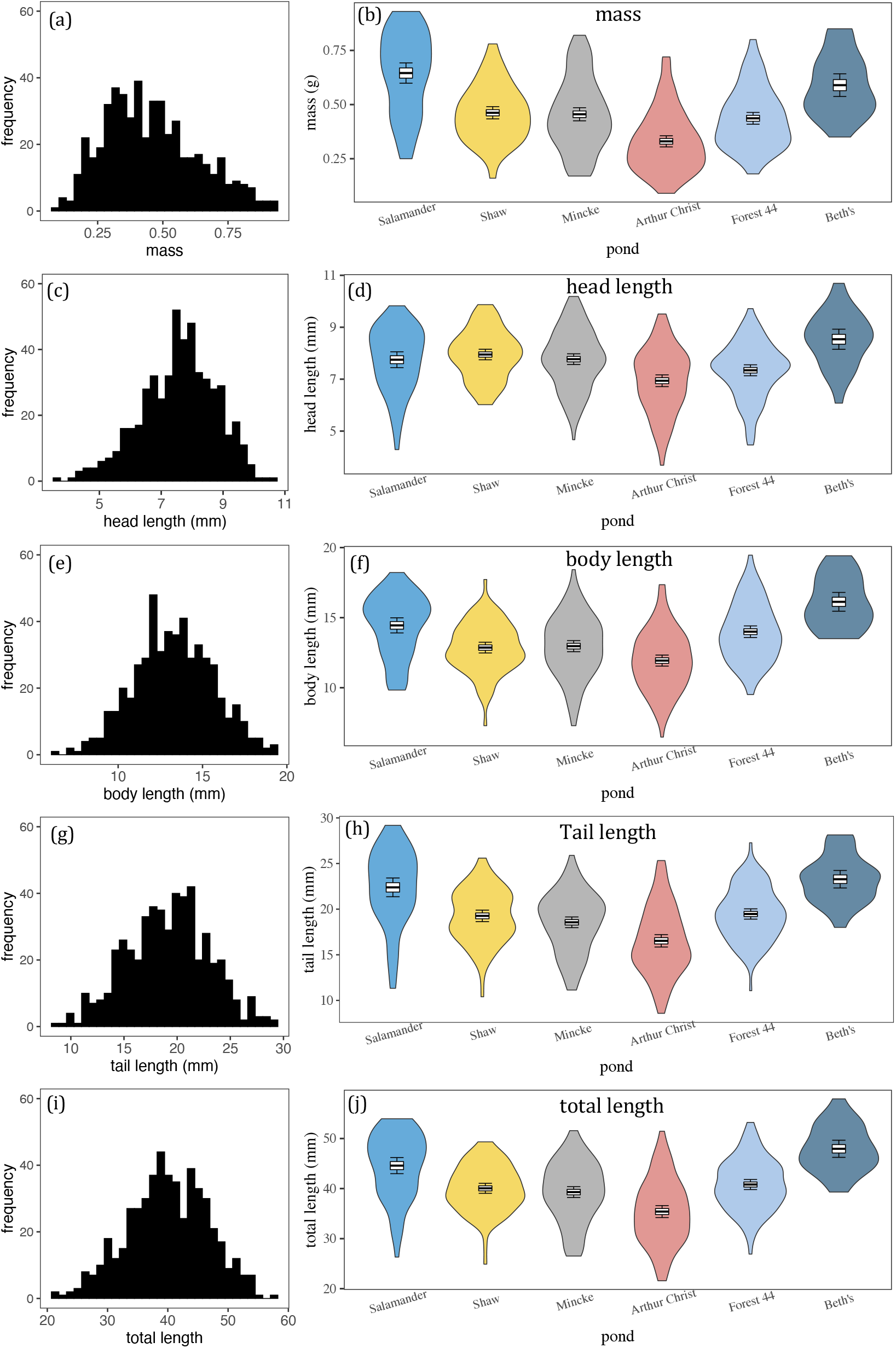
Salamander mass and length variation within and between ponds. Panels on left **(a,c,e,g,i)** are frequency distributions of mass and length measurements across all focal ponds. Violin plots on the right-side panels **(b,d,f,h,j)** distinguish individual-level and between-pond morphological variation to show how the traits were spatially structured. Box plots within the violin plots denote the mean, standard error, and 95% confidence intervals of trait measures. Ponds in the right-side panels are ordered from left to right in the chronological order in which they were sampled.

We then assessed within- and between-pond variation in mass-length allometry, a form of trait co-variation. Specifically, we ran GLMMs to test whether the slopes and intercepts of mass-length regressions differed across the six focal ponds, again distinguishing head length, body length, and tail length. GLMMs included mass as the response, length as a fixed effect, pond as a random intercept term, and length as a random slope term. We log-transformed both length and mass and used a Gaussian error structure for the normalized data in all models. To enable convergence of these more complex models, we multiplied (log-transformed) mass by a factor of 10 to standardize the units with length measurements. To test for differences in regression intercepts and slopes across focal ponds, we used likelihood ratio tests comparing the fit of models that included both terms with models omitting the random intercept or random slope term. We also calculated the marginal and conditional R^2^ of the models using the *MUMIn* package in R. Marginal R^2^ is a measure of the amount of variation in mass that was explained by the fixed effect of length, while conditional R^2^ considers variation explained by both fixed and random effect terms (Johnson, 2014).

To further assess differences in the co-variation of morphological traits among focal ponds, we determined the extent to which between-pond variation in salamander head, body, tail, and overall (all segments combined) shape contributed to total observed variation in these multidimensional morphological traits. Again, we calculated the ICC from GLMMs, including ‘pond’ as a random intercept term. PC1 scores were used as response variables. The shape data were also normally distributed (Fig. 3), so we used a Gaussian error structure for all models.

To perform our preliminary test of the influence of pond age and predator density on salamander mass, length, and shape, we ran GLMMs that included pond age (new vs. old) and predator density as fixed effects, and the pond name as a random effect. To test how the age and predator density of ponds influenced mass-length co-variation, we ran GLMMs with log-transformed mass as the response and log-transformed length, the focal factor (i.e., pond age or predator density) and their interaction as fixed effects. We also included salamander length (again, log-transformed) as a random slope term, and pond name as a random intercept term. We ran separate GLMMs for the two interactions to prevent model overfitting. We also ran separate GLMMs for our different length measures: head length, body length, tail length, and total length. We compared the fit of models including the interaction terms with models omitting the interaction terms, using likelihood ratio tests, to test the influence of pond age and predator density on the relationship between mass and length.

## Results

We measured a total of 519 spotted salamander larvae (Forest 44: N = 101, Shaw: N = 88, Salamander pond: N = 65, Arthur Christ: N = 116, Beth’s: N = 30, Mincke: N = 119) in this study. Salamander mass was not influenced by their feeding on the previous day in the subset of 227 individuals tested (*F*_(1-82)_ = 3.43, *p* = 0.068). Within all ponds, salamander mass varied more among individuals than any of the length measurements (Fig. 2a). Morphological variation was consistently lower in Beth’s Pond (Fig. 2b), although this was likely due to the lower sample size. Otherwise, there was no indication that specific ponds had more or less morphological variation (Figs. 2b and 3).

Between-pond differences in average trait values accounted for 7 to 35% of the total observed variation in salamander mass and length, depending on the specific body section measured (Table 1). Specifically, between-pond differences accounted for proportionally more of the observed variation in salamander mass (35%), total length (35%), and tail length (27%) than in head (11%) and body length (17%) (Table 1).

**Table 1.** Proportion of mass, length, and shape variation attributable to between-pond differences. Displayed are the variance components of generalized linear mixed models used for our analyses of within- and between-pond variation in salamander mass, length, and shape. The intra-class correlation coefficient (ICC), or the variance partitioning component, is the proportion of between-pond variation explaining total observed trait variation (between-pond variance + residual variance). Higher ICC values denote higher degree of between-pond variation in salamander traits. CI = confidence intervals of the ICC derived from bootstrapping over 500 resampling events.

| body segment | between-pond variance | SD | residual variance | SD | ICC (%) | 95% CI - lower | 95% CI - upper |
| --- | --- | --- | --- | --- | --- | --- | --- |
| **mass** |  |  |  |  |  |  |  |
|  | 0.013 | 0.112 | 0.023 | 0.152 | 35.3 | 8.62 | 58.71 |
| **length** |  |  |  |  |  |  |  |
| head | 0.15 | 0.39 | 1.27 | 1.13 | 10.8 | 0.73 | 26.11 |
| body | 0.92 | 0.96 | 4.36 | 2.09 | 17.4 | 8.72 | 53.62 |
| tail | 4.27 | 2.07 | 11.29 | 3.36 | 27.5 | 4.08 | 52.07 |
| combined | 18.23 | 4.27 | 32.91 | 5.74 | 35.6 | 6.25 | 57.80 |
| **shape** |  |  |  |  |  |  |  |
|  | between-pond variance | SD | residual variance | SD | ICC | 95% CI - lower | 95% CI - upper |
| head | 0.0004 | 0.019 | 0.003 | 0.053 | 11.0 | 0.7 | 26.5 |
| body | 0.0002 | 0.016 | 0.002 | 0.049 | 9.4 | 0.3 | 23.6 |
| tail | 0.001 | 0.026 | 0.002 | 0.049 | 22.8 | 1.2 | 35.0 |
| combined | 0.000 | 0.009 | 0.001 | 0.033 | 7.60 | 0.2 | 17.7 |

Salamander mass strongly co-varied with total length, which makes sense because we took single measures of mass that incorporated all body segments. Mass also strongly co-varied with body length and tail length, but was less correlated with head length, likely because heads are the smallest body segment of salamanders (Fig. S1). There were detectable differences in the intercepts of mass-length relationships across ponds (including pond as a random intercept term improved model fit; mass-head length: *X*^2^_1_ = 111.84, p < 0.001; mass-body length: *X*^2^_1_ = 141.09, p < 0.001; mass-tail length: *X*^2^_2_ = 39.089, p < 0.001; total length: *X*^2^_1_ = 81.136, p < 0.001; Fig. 4). For a given length, individuals from Salamander Pond tended to be heavier than individuals from other ponds (Fig. 4) whereas individuals form Beth’s Pond were generally lighter in mass per unit length (Fig. 4). The slopes of mass-length relationships were less influenced by pond of capture (Table 2). Only the slopes for mass-head length and mass-tail length relationships were influenced by the pond of capture (mass-head length: *X*^2^_2_ = 8.68, p = 0.013; mass-tail length: *X*^2^_2_ = 11.25, p = 0.004; Fig. 4); the slopes of mass-body length and mass-total length relationships were consistent across sampled ponds (mass-body length: *X*^2^_2_ 4.24, p = 0.120; mass-total length: *X*^2^_2_ = 5.05, p = 0.080; Fig. 3). The slope exponents were always < 3 (Table S3), indicating that larger salamander larvae generally had more elongate heads, bodies, and tails than smaller larvae.

**Fig. 4.**
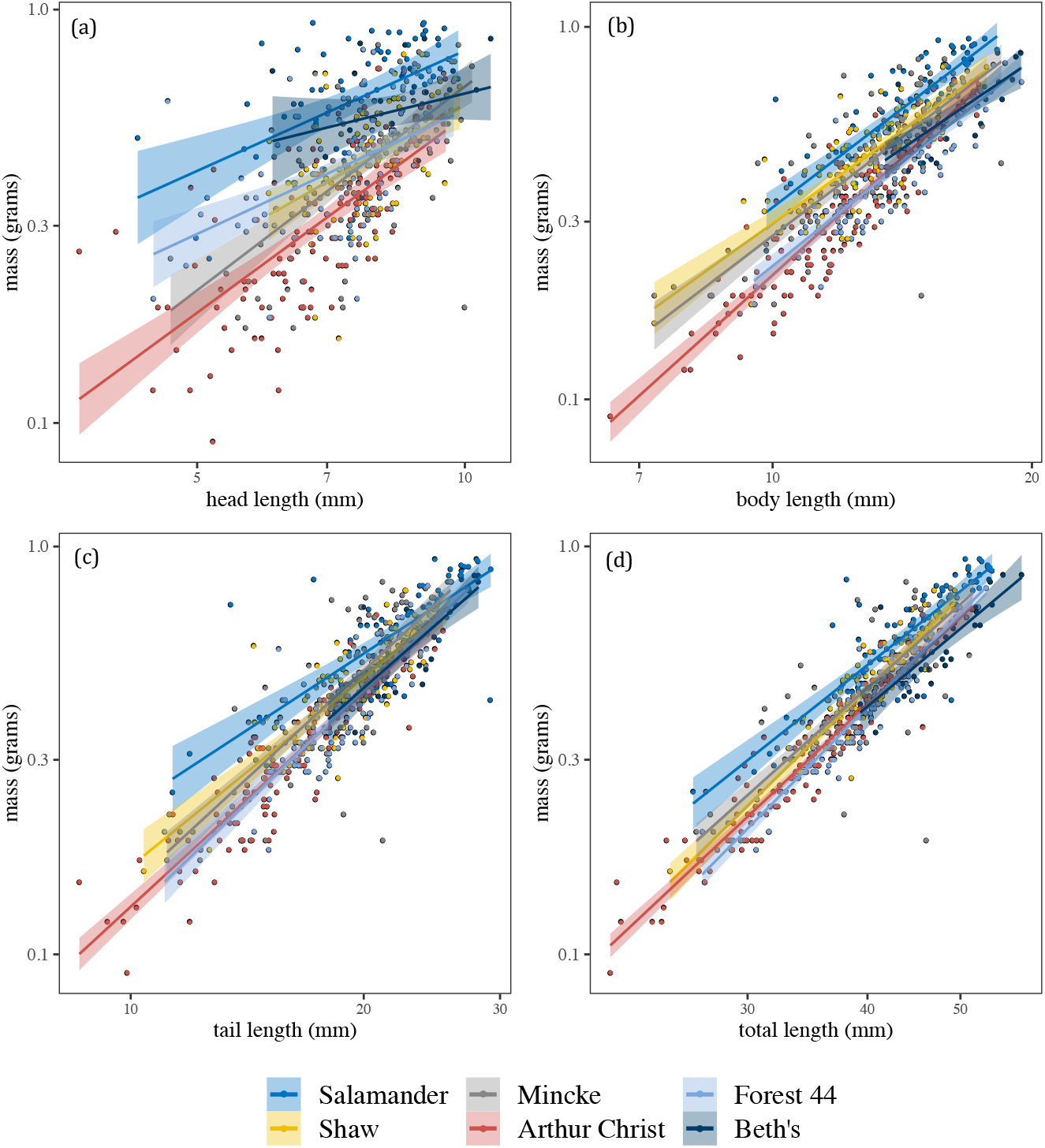
Between-pond variation in mass-length allometry in salamanders. Regression lines of salamander mass with (a) head length, (b) body length, (c) tail length, and (d) are shown for the six focal ponds, distinguished by line colors. Shaded areas show the 95% confidence intervals of the regression lines. Mass and length are plotted on a log_10_ scale in all cases.

**Table 2.** Variation in intercepts and slopes of mass-length relationships. Displayed are the variance components of generalized linear mixed models used for our analyses of between-pond variation in the scaling of salamander mass with different length measurements. Models included length (log-transformed) as a fixed effect and a random slope term, and pond as a random intercept term. Marginal R^2^ denotes the amount of variation in mass explained by the length measurement alone, whereas conditional R^2^ considers variation explained by the random intercept and slope terms.

| mass-length co-variation |  |  |  |  |  |  |  |  |
| --- | --- | --- | --- | --- | --- | --- | --- | --- |
| body segment | intercept variance | SD | slope variance | SD | residual variance | SD | marginal R<sup>2</sup> | conditional R<sup>2</sup> |
| head | 0.11 | 0.33 | 0.09 | 0.29 | 0.02 | 0.13 | 0.26 | 0.49 |
| body | 0.04 | 0.20 | 0.02 | 0.13 | 0.01 | 0.09 | 0.65 | 0.77 |
| tail | 0.08 | 0.28 | 0.04 | 0.19 | 0.01 | 0.08 | 0.73 | 0.79 |
| combined | 0.07 | 0.26 | 0.00 | 0.07 | 0.00 | 0.07 | 0.81 | 0.87 |

Similar to length measures, the shape of salamander tails exhibited more between-pond variation than did the heads or bodies. Between-pond trait differences contributed 25% of the total observed variation in tail shape, compared with 11%, 9%, and 7% of head, body, and overall shape, respectively (Table 1). Furthermore, there was little evidence that PC scores were clustered by pond (Fig. 5), indicating a weak signal of between-pond variation in salamander shape.

**Fig. 5.**
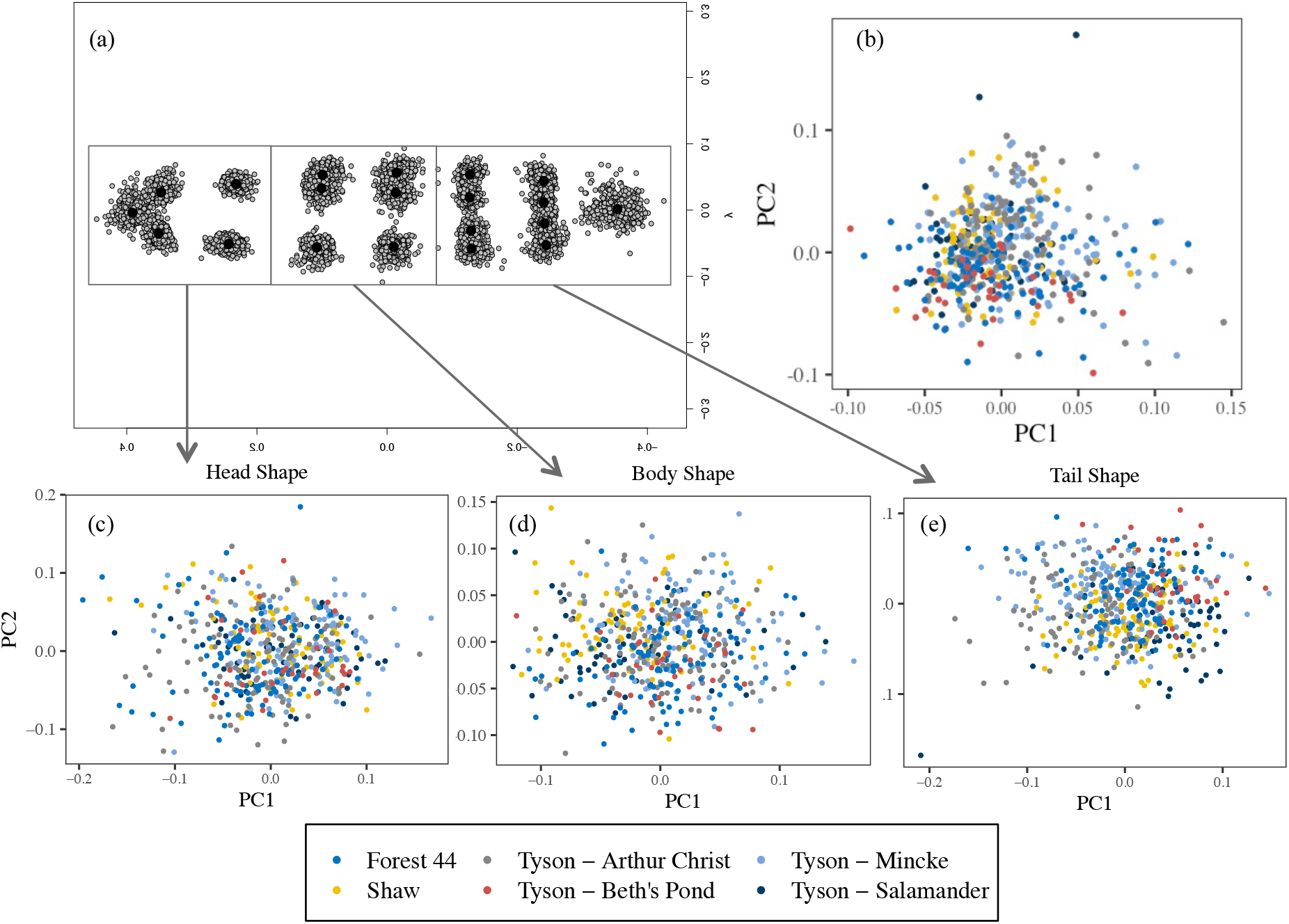
Salamander shape variation within and among ponds. The shape values are based on sets of landmarks at different points along the lateral surface of salamander bodies (a). The overall (b), head (c), body (d), and tail (e) shape of salamanders collected from the six focal ponds. The shape values are based on sets of landmarks at different points along the lateral surface of salamander bodies. PC1 and PC2 values increase with elongation of shape and increasing length:height ratio.

Neither pond age nor predator density influenced salamander mass or any measures of length (Table S4). However, these pond attributes did influence certain mass-length relationships and body shapes (Fig. S4, Table S5). Pond age influenced the scaling of mass with head and tail length (Fig. S4, Table S5). In contrast, predator density influenced mass-body length relationships (Fig S2, Table S2). Both pond age and predator density influenced salamander head shape, but neither influenced body or tail shape. Pond age also influenced the overall shape of salamanders, but predator density did not (Table S4).

## Discussion

Measuring the lengths and masses of salamander larvae in a network of spatially discrete ponds showed that most morphological variation and co-variation occurred within ponds. Between-pond morphological differences were not negligible however, particularly for salamander tails. Salamander tails exhibited more between-pond variation than heads or bodies. Between-pond differences in trait co-variation were also evident in salamander tails. Scaling of mass with salamander tail lengths differed across ponds both in terms of the intercepts and the slopes of the relationships, and between-pond differences contributed more to total observed variation in tail shape than for head or body shape. The spatial discreteness of pond habitats therefore seems to act strongly on salamander tail morphology (see below for further discussion).

The substantial within-pond variation in salamander morphology suggests that local factors, such as microhabitat heterogeneity, influence salamander morphology. Given that many ponds were spaced within the documented dispersal ranges of salamanders (Zamudio and Wieczorek, 2006; Patrick et al., 2008), movement between ponds could also have reduced the contribution of between-pond differences to morphological variation by sustaining mixing of genotypes and phenotypes. At the metapopulation scale, local and spatial factors likely interact to shape varying degrees of between-pond morphological variation similar to what we observed in salamanders.

The stronger contribution of between-pond differences to salamander tail variation may be because tails play an important role in locomotive (i.e., swimming) performance. Being meso-predators, swimming performance for salamanders is critical to both capturing prey and evading predators (Van Buskirk and Schmidt, 2000; Urban, 2010; Landberg and Azizi, 2010). Tails may therefore be more closely linked to fitness, hence under stronger selection, than heads and bodies, at least in habitats where predation is a significant threat (Landberg and Azizi, 2010). Multiple predators of larval salamanders were found in our focal ponds, so predation risk is likely to be a strong selective force in the salamander metapopulation studied here. In a preliminary analysis of the data, we did not detect an influence of predator density on tail length of salamanders (see Supplementary Material), but this analysis was based on our limited sample of ponds and warrants further investigation. Regardless of the factors driving salamander tail variation, our findings suggest that salamander responses to habitat alterations, biological invasions, and other pond-level disturbances may manifest as changes in tail morphology as opposed to changes in head and body morphology. If this prediction holds, the more common body size measure for amphibians, snout-to-vent length, which only takes head and body length into account, would be insufficient for predicting the eco-evolutionary responses of this species to landscape-scale environmental changes in aquatic habitats.

Between-pond variation in the allometric relationship between salamander mass and total length arose specifically from differences in intercepts; slopes of the relationship were highly consistent. Pond-level effects appear to act on the relationship of salamander mass to total body length, but they do not appear to alter how mass scales with total body length. This spatially robust scaling of mass and total body length may explain why the two traits separately exhibited nearly identical degrees of between-pond variation. More broadly, this pattern of allometric scaling, in which intercepts but not slopes of trait relationships differ, is consistent with allometric relationships documented across many other taxa (Voje et al., 2014), suggesting a general constraint to the plasticity and evolution of the slopes of trait relationships.

One important caveat to the above findings is that our sampling could not distinguish between-pond variation in salamander morphology from possible temporal variation in salamander morphology. Spotted salamander larvae exhibit growth and developmental changes within summer months that could have contributed to observed morphological variation. We expect the contributions of temporal changes to salamander morphological variation to have been minor because we sampled salamander larvae during latter developmental stages, evidenced by all salamanders being in the final Harrison stages (45-46) (Harrison, 1969) after most growth and development had occurred (Landberg and Azizi, 2010). In support of this expectation, body lengths and mass did not increase monotonically throughout the sampling period, which should be the case when growth and development drive morphological variation. Nevertheless, we cannot rule out the possible influence of temporal morphological variation and advocate for further work that corroborates our findings through longitudinal pond sampling, which would account for growth and ontogeny.

The inclusion of morphological diversity data in biodiversity conservation stems from the idea that different populations of the same species are not equal in terms of eco-evolutionary history. As such, exploring various approaches to the conservation of morphological diversity is important to developing strategies for reducing biodiversity losses under global change (Des Roches et al., 2018). The mix of within- and between-pond morphological variation in salamanders provides promise that pond construction can utilize local and spatial processes to bolster morphological diversity. Capitalizing on the presence of new constructed ponds in our study area, we made a preliminary comparison of salamander morphology and allometry between new and old ponds (see Supplementary Material for detailed methods and results). Although our analysis did not detect differences in mass or length of salamanders between new and old ponds, we did find differences in mass-length relationships and body shape (Supplementary Material Table S4). Habitat restoration through pond construction may therefore bolster diversity in trait co-variation, and in so doing may instill more adaptive potential under environmental change (Laughlin and Messier, 2015). Although there are several studies for various taxa that quantify functional connectivity between habitat patches (and local populations) using genetic techniques, we encourage additional studies on morphological parameters and patterns to better understand the mechanisms that promote long-term population persistence in fragmented landscapes.

## Acknowledgements

We thank members of the Tyson Research Station for their support of our field sampling, J. Grady for assistance with data analyses and visualization, J. Messier for assistance with the statistical analyses, and S. Trombulak for assistance with data analyses and comments that significantly improved this manuscript. This project was conducted in accordance with University of Illinois IACUC #16203 and funded, in part, by the NGRREC intern program. None of the authors experienced conflict of interest that could have influenced the objectivity of this study.

## Author Contributions

DRD and ETG designed the study and executed field sampling and trait measurements, with supervisorial support from AID and JAC. ETG completed the morphometrics and principal components analyses. DRD performed the rest of the statistical analyses. All authors contributed to the writing and revision of the manuscript.

