## Supplementary Material for "Spatial context of trait variation: morphology of spotted salamanders (*Ambystoma maculatum*) varies more within ponds than between ponds"

### Effect of salamander feeding on mass

All salamanders were fed one tree frog tadpole the day prior to being weighed, but not all salamanders ate the frogs. Acknowledging that differential feeding may influence our mass measurements, we statistically tested whether mass of salamanders that completely consumed tadpoles differed from salamanders that did not. Specifically, we ran generalized linear models with mass as the response and feeding outcome as a categorical fixed effect (consumed or not consumed). Models had a Gaussian error structure because mass values were normally distributed (Fig. 3 in main text). We tested the influence of feeding outcome on mass by comparing a model with feeding outcome as a fixed effect with a model omitting the effect, using a likelihood ratio test. We used a random subset of 237 salamanders for which we had feeding records. Including feeding outcome in the model did not significantly improve model fit ($F_{(1\text{-}82)} = 3.43$, $p = 0.068$). Thus, we did not include feeding outcome as a factor in models used in the main analyses.

### Tables and Figures

| Pond Name | sample size | Age | predator Density | Diameter (m) |
|---|---|---|---|---|
| Forest 44 | 105 | old | 3.1 | 30 |
| Shaw | 90 | old | 1.9 | 48 |
| Salamander | 69 | old | 0.3 | 69 |
| Arthur Christ | 117 | new | 11.5 | 8 |
| Beth’s | 30 | new | 10 | 8 |
| Mincke | 118 | new | 2 | 8 |

**Table S1. Pond characteristics.** Environmental and ecological variables of the six ponds in east-central Missouri from which salamander larvae were collected.

| trait | pond | mean | sd | min | max | CV |
|---|---|---|---|---|---|---|
| **mass (g)** | Salamander | 0.65 | 0.19 | 0.25 | 0.93 | 29.29 |
| | Shaw | 0.46 | 0.13 | 0.16 | 0.78 | 28.39 |
| | Mincke | 0.46 | 0.17 | 0.17 | 0.82 | 36.65 |
| | Arthur Christ | 0.33 | 0.14 | 0.09 | 0.72 | 42.43 |
| | Forest 44 | 0.44 | 0.14 | 0.18 | 0.80 | 31.59 |
| | Beth's | 0.59 | 0.14 | 0.35 | 0.85 | 23.79 |
| **head length (mm)** | Salamander | 7.75 | 1.23 | 4.29 | 9.83 | 15.88 |
| | Shaw | 7.95 | 0.94 | 6.02 | 9.87 | 11.84 |
| | Mincke | 7.77 | 1.14 | 4.67 | 10.19 | 14.68 |
| | Arthur Christ | 6.94 | 1.23 | 3.69 | 9.51 | 17.78 |
| | Forest 44 | 7.34 | 1.07 | 4.47 | 9.72 | 14.57 |
| | Beth's | 8.54 | 1.04 | 6.08 | 10.69 | 12.17 |
| **body length (mm)** | Salamander | 14.45 | 2.18 | 9.84 | 18.22 | 15.08 |
| | Shaw | 12.86 | 1.78 | 7.29 | 17.72 | 13.81 |
| | Mincke | 12.97 | 2.18 | 7.29 | 18.43 | 16.82 |
| | Arthur Christ | 11.94 | 2.14 | 6.48 | 17.35 | 17.97 |
| | Forest 44 | 14.00 | 2.10 | 9.51 | 19.46 | 14.98 |
| | Beth's | 16.13 | 1.79 | 13.50 | 19.42 | 11.12 |
| **tail length (mm)** | Salamander | 22.39 | 4.14 | 11.33 | 29.18 | 18.47 |
| | Shaw | 19.26 | 2.92 | 10.40 | 25.60 | 15.16 |
| | Mincke | 18.56 | 3.24 | 11.14 | 25.90 | 17.43 |
| | Arthur Christ | 16.53 | 3.69 | 8.58 | 25.32 | 22.35 |
| | Forest 44 | 19.47 | 2.86 | 11.06 | 27.27 | 14.70 |
| | Beth's | 23.28 | 2.57 | 18.00 | 28.14 | 11.04 |
| **total length (mm)** | Salamander | 44.59 | 6.49 | 26.32 | 53.94 | 14.56 |
| | Shaw | 40.05 | 4.64 | 24.90 | 49.33 | 11.58 |
| | Mincke | 39.30 | 5.98 | 26.53 | 51.57 | 15.22 |
| | Arthur Christ | 35.41 | 6.50 | 21.58 | 51.45 | 18.35 |
| | Forest 44 | 40.81 | 5.10 | 26.91 | 53.21 | 12.49 |
| | Beth's | 47.95 | 4.57 | 39.31 | 57.92 | 9.54 |

**Table S2. Summary of salamander length (head, body, tail, total) and mass.** Summary statistics of 2016 survey data of late-phase larval salamander populations in Missouri. Sd – standard deviation, min = minimum value, max = maximum value, CV = coefficient of variation.

| Trait | Pond name | mass regression equation |
|---|---|---|
| **head length** | Forest 44 | y = 1.00x -1.24 |
| | Shaw | y = 1.21x -1.44 |
| | Salamander | y = 0.97x -1.07 |
| | Arthur Christ | y = 1.58x -1.84 |
| | Beth's | y = 0.53x -0.74 |
| | Mincke | y = 1.64x -1.82 |
| | **Overall** | **y = 1.52x -1.71** |
| **body length** | Forest 44 | y = 1.84x -2.49 |
| | Shaw | y = 1.67x -2.2 |
| | Salamander | y = 1.76x -2.25 |
| | Arthur Christ | y = 2.09x -2.76 |
| | Beth's | y = 1.57x -2.13 |
| | Mincke | y = 1.76x -2.32 |
| | **Overall** | **y = 1.94x -2.54** |
| **tail length** | Forest 44 | y = 1.83x -2.73 |
| | Shaw | y = 1.52x -2.3 |
| | Salamander | y = 1.24x -1.88 |
| | Arthur Christ | y = 1.75x -2.63 |
| | Beth's | y = 1.67x -2.52 |
| | Mincke | y = 1.75x -2.59 |
| | **Overall** | **y = 1.74x -2.6** |
| **total length** | Forest 44 | y = 2.38x -4.21 |
| | Shaw | y = 2.3x -4.04 |
| | Salamander | y = 1.87x -3.28 |
| | Arthur Christ | y = 2.21x -3.92 |
| | Beth's | y = 1.96x -3.55 |
| | Mincke | y = 2.11x -3.73 |
| | **Overall** | **y = 2.23x -9.94** |

**Table S3. Length-mass regression equations for salamander morphology.** Equations for regression lines expressing the relationship between mass with the length of salamander heads, bodies, tails, and the three body segments combined (total length). Both length and mass values were log-transformed in linear models used to calculate intercept and slope values for the regression lines.

| body segment | Factor | df | AIC | $X^2$ | p |
|---|---|---|---|---|---|
| **mass** | | | | | |
| | pond age | 1 | -452.73 | 0.00 | 0.949 |
| | predator density | 1 | -452.29 | 0.45 | 0.505 |
| **length** | | | | | |
| head | pond age | 1 | 1617.20 | 0.26 | 0.613 |
| | predator density | 1 | 1617.30 | 0.37 | 0.541 |
| body | pond age | 1 | 2245.60 | 0.05 | 0.816 |
| | predator density | 1 | 2245.60 | 0.04 | 0.841 |
| tail | pond age | 1 | 2732.60 | 0.10 | 0.757 |
| | predator density | 1 | 2732.50 | 0.01 | 0.909 |
| total | pond age | 1 | 3296.80 | 0.04 | 0.848 |
| | predator density | 1 | 3296.80 | 0.00 | 0.950 |
| **mass:length co-variation** | | | | | |
| head | pond age | **1** | **-588.61** | **12.21** | **0.002** |
| | predator density | 1 | -624.45 | 2.87 | 0.091 |
| body | pond age | 1 | -1030.80 | 1.94 | 0.163 |
| | predator density | **1** | **-1034.30** | **6.49** | **0.011** |
| tail | pond age | **1** | **-1073.50** | **5.43** | **0.020** |
| | predator density | 1 | -1078.30 | 3.44 | 0.064 |
| total | pond age | 1 | -1296.80 | 0.13 | 0.717 |
| | predator density | 1 | -1300.90 | 0.39 | 0.534 |
| **shape** | | | | | |
| head | **pond age** | **1** | **-1550.50** | **5.68** | **0.017** |
| | **predator density** | **1** | **-1549.80** | **6.40** | **0.011** |
| body | pond age | 1 | -1647.20 | 2.82 | 0.093 |
| | predator density | 1 | -1649.50 | 0.55 | 0.459 |
| tail | pond age | 1 | -1633.80 | 0.80 | 0.372 |
| | predator density | 1 | -1634.30 | 0.26 | 0.612 |
| overall | pond age | **1** | **-2052.60** | **5.53** | **0.019** |
| | predator density | 1 | -2056.30 | 1.78 | 0.182 |

**Table S4. Influence of pond age and predator density on salamander morphology.** Outputs of likelihood ratio tests of the influence of pond age and predator density on salamander morphological traits are reported. For mass-length co-variation, we tested the effects of pond age and predator density with separate models, to enable convergence of the complex models. Df = degrees of freedom, AIC = Akaike's Information Criterion. Cases where the factors significantly improved model fit to the data are highlighted in bold.

| Invertebrate Species | Predatory (Y/N) | Beth's pond | Mincke pond | Salamander pond | Shaw pond | Arthur Christ pond | Forest 44 pond | total |
|---|---|---|---|---|---|---|---|---|
| *Acilius fraternus* | yes | 0 | 0 | 1 | 0 | 0 | 0 | 1 |
| Acilius sp. larvae | yes | 0 | 3 | 0 | 0 | 0 | 0 | 3 |
| *Aeshna umbrosa* | yes | 0 | 0 | 1 | 0 | 0 | 0 | 1 |
| Agabus sp. larvae | yes | 2 | 1 | 0 | 0 | 0 | 0 | 3 |
| *Anaxyrus americanus* | no | 114 | 0 | 0 | 0 | 0 | 0 | 114 |
| Chaoborus sp. larvae | no | 108 | 38 | 0 | 0 | 0 | 0 | 146 |
| Chauliodes sp. larvae | yes | 0 | 0 | 5 | 0 | 0 | 0 | 5 |
| Chironomid sp. | no | 89 | 153 | 0 | 0 | 0 | 0 | 242 |
| Enallagma sp. | no | 1 | 0 | 0 | 0 | 1 | 0 | 2 |
| *Erythemis simplicollis* | yes | 0 | 0 | 0 | 0 | 5 | 0 | 5 |
| *Gyraulus parvus* | no | 0 | 3 | 0 | 0 | 0 | 0 | 3 |
| *Helisoma trivolvis* | no | 0 | 2 | 0 | 0 | 0 | 0 | 2 |
| Helobdella sp. | no | 0 | 7 | 0 | 0 | 0 | 0 | 7 |
| Hesperocorixa sp. | no | 0 | 0 | 1 | 0 | 0 | 0 | 1 |
| Hydrobiomorpha sp. | no | 0 | 0 | 1 | 0 | 0 | 0 | 1 |
| *Hyla versicolor* | no | 0 | 4 | 0 | 0 | 0 | 0 | 4 |
| *Laccophilus maculosa* | no | 1 | 2 | 0 | 0 | 0 | 1 | 4 |
| Laccophilus sp. larvae | no | 0 | 2 | 0 | 0 | 0 | 0 | 2 |
| *Libellula luctuosa* | yes | 7 | 0 | 0 | 0 | 0 | 0 | 7 |
| *Libellula pulchella* | yes | 4 | 0 | 0 | 0 | 0 | 0 | 4 |
| *Musculium transversum* | no | 0 | 200 | 0 | 0 | 0 | 0 | 200 |
| *Notonecta irrorata* | yes | 0 | 0 | 0 | 3 | 3 | 14 | 20 |
| *Notophthalamus viridescens louisianensis* | yes | 1 | 0 | 0 | 0 | 0 | 0 | 1 |
| Ogliochaete | no | 0 | 2 | 0 | 0 | 0 | 0 | 2 |
| *Pachydiplax longipennis* | yes | 5 | 0 | 0 | 2 | 15 | 0 | 22 |
| *Physa heterostropha* | no | 57 | 13 | 0 | 0 | 0 | 0 | 70 |
| *Pseudacris triseriata* | no | 0 | 4 | 0 | 0 | 0 | 0 | 4 |

| | | | | | | | | |
|---|---|---|---|---|---|---|---|---|
| Pseudosuccinea sp. | no | 1 | 0 | 0 | 0 | 0 | 0 | 1 |
| *Rana clamitans* | yes | 3 | 0 | 0 | 25 | 0 | 0 | 28 |
| *Tropisternus blachelyi* | yes | 0 | 0 | 0 | 0 | 0 | 16 | 16 |
| Tropisternus sp. 1 | yes | 0 | 0 | 0 | 0 | 0 | 1 | 1 |
| Tropisternus sp. 4 | yes | 0 | 0 | 1 | 1 | 0 | 0 | 2 |
| Tropisternus sp. larvae | no | 0 | 0 | 0 | 0 | 0 | 3 | 3 |

**Table S5.** Counts of different species of invertebrates of focal ponds where salamanders were collected. Counts were performed within the same time period – July-August 2016 - as when salamander collected was executed, except for Beth's pond. Counts for Beth's pond come from a 2013 survey.

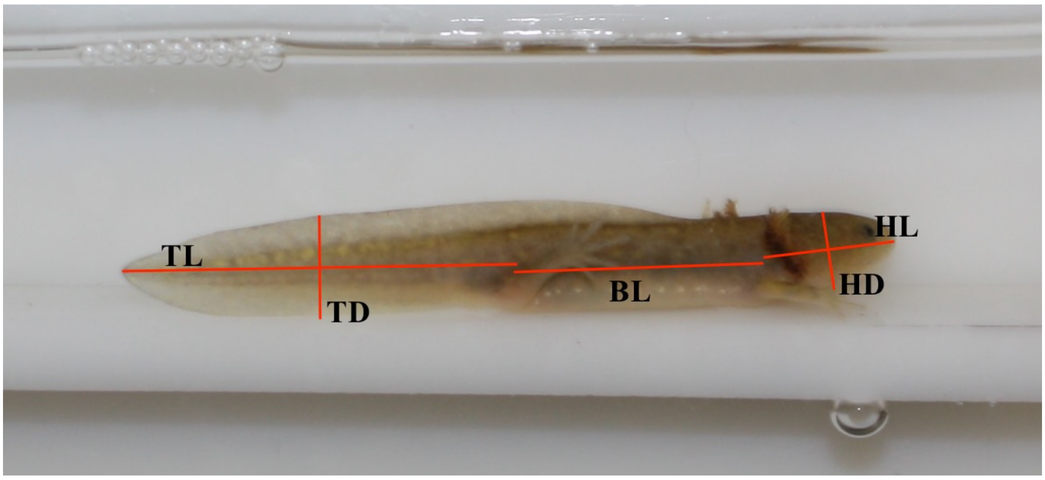


**(b)**

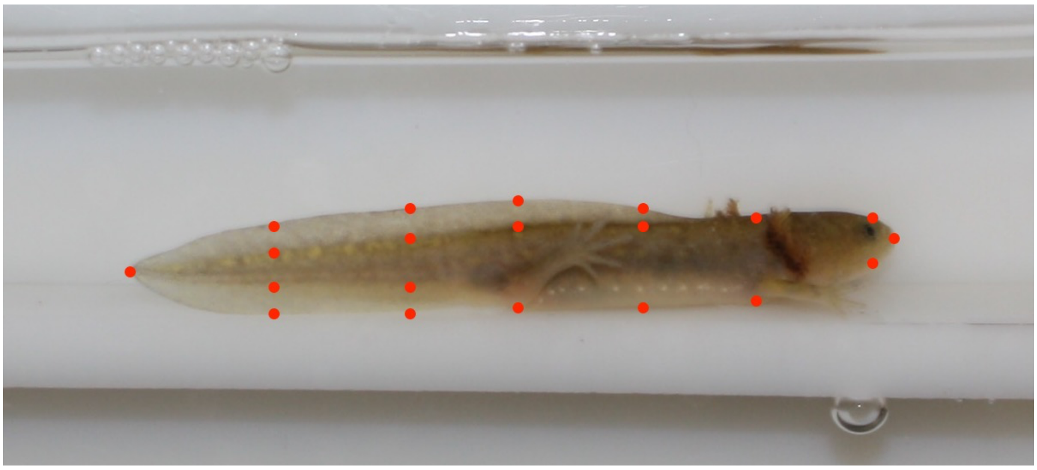

**Fig. S1.** Morphometric analysis of larval *A. maculatum* included both **(a)** measurements and **(b)** landmarks placed for shape analysis. Linear measurements included head length (HL), maximum head depth (HD), body length (BL), tail length (TL), and mid tail depth (TD). Geometric analysis involved placing twenty landmarks to outline the shape of, and included: tip of the snout (1) above the eye (2), below the eye (3), above vent (4), below vent (5), 50% length of body (6, 7, 8), in line with hind leg (9, 10, 11), at 25% length of tail (12, 13, 14, 15), at 50% length of tail (16, 17, 18, 19), and tip of tail (20).

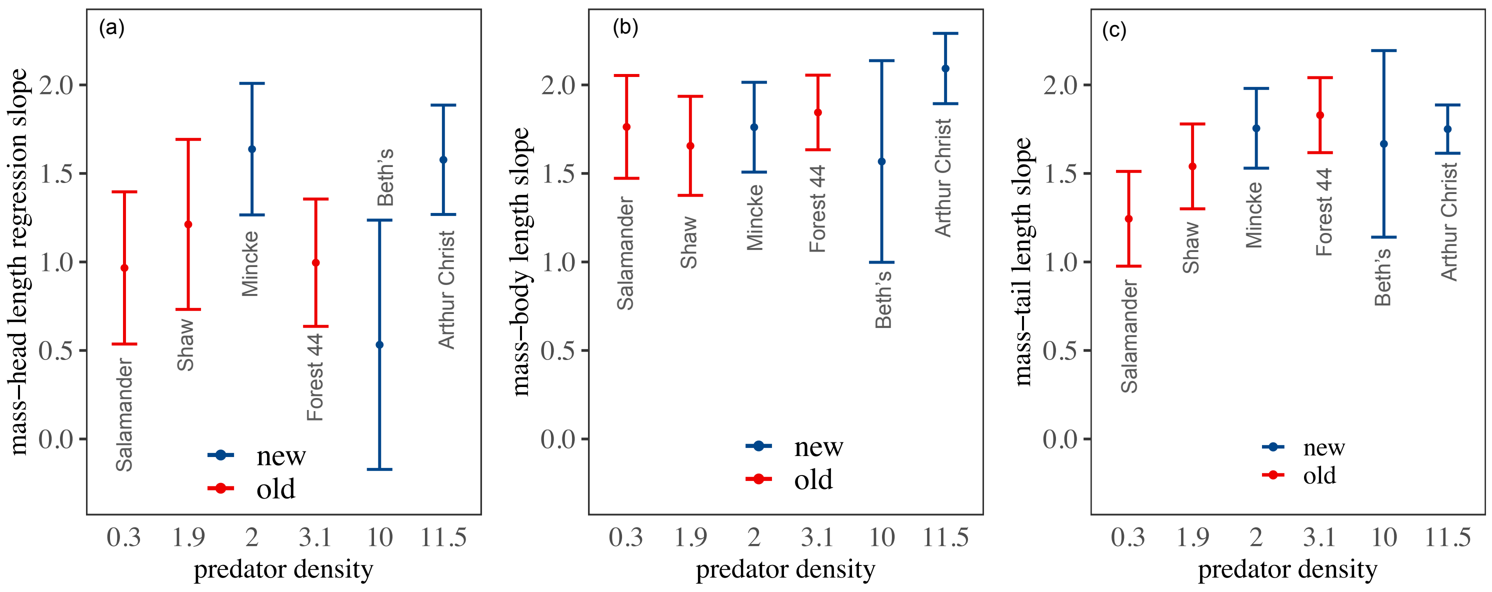


**Fig. S2. The effect of pond age and predator density on mass-length relationships.** The slope values for co-variation of salamander mass with (a) head length, (b) body length, and (c) tail length are displayed according to the predator density in ponds. Predators comprised invertebrates and adult newts. Red bars denote 'old' ponds that were constructed in the mid-1900s, and blue bars denote 'new' ponds that were constructed in 2008. Error bars denote 95% confidence intervals.

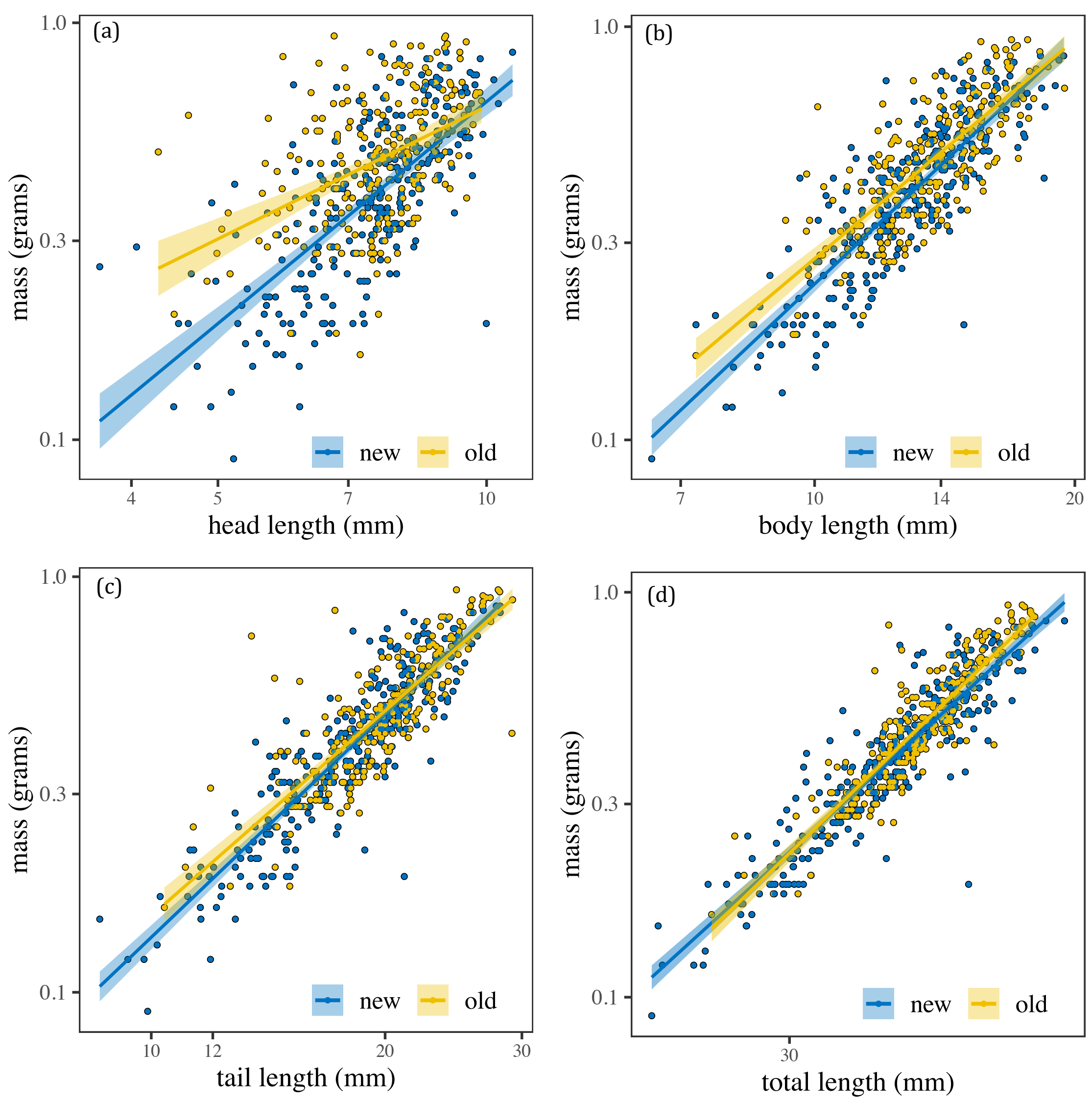


**Fig. S3. The effect of pond age on mass-length relationships.** Scaling of salamander mass with (a) head length, (b) body length, (c) tail length, and (d) total length are displayed, with regression lines drawn separately for new (blue) and old (yellow) ponds. New ponds were constructed in 2008. Old ponds were constructed in the mid-1900s. Shaded areas denote 95% confidence intervals.
